# Predicting ligand-dependent tumors from multi-dimensional signaling features

**DOI:** 10.1101/142901

**Authors:** Helge Hass, Kristina Masson, Sibylle Wohlgemuth, Violette Paragas, John E Allen, Mark Sevecka, Emily Pace, Jens Timmer, Joerg Stelling, Gavin MacBeath, Birgit Schoeberl, Andreas Raue

## Abstract

Targeted therapies have shown significant patient benefit in about 5-10% of solid tumors that are addicted to a single oncogene. Here, we explore the idea of ligand addiction as a driver of tumor growth. High ligand levels in tumors have been shown to be associated with impaired patient survival, but targeted therapies have not yet shown great benefit in unselected patient populations. Using a novel approach of applying Bagged Decision Trees (BDT) to high-dimensional signaling features derived from a computational model, we can predict ligand dependent proliferation across a set of 58 cell lines. This mechanistic, multi-pathway model that features receptor heterodimerization, was trained on seven cancer cell lines and can predict signaling across two independent cell lines by adjusting only the receptor expression levels for each cell line. Interestingly, for patient samples the predicted tumor growth response correlates with high growth factor expression in the tumor microenvironment, which argues for a co-evolution of both factors *in vivo*.

**Summary:** Prediction of ligand-induced growth of cancer cell lines, which correlates with ligand-blocking antibody efficacy, could be significantly improved by learning from features of a mechanistic signaling model, and was applied to reveal a correlation between growth factor expression and predicted response in patient samples.

## Introduction

The combination of Herceptin® with chemotherapy demonstrated a dramatically increased survival benefit for a subset of women with HER2 amplified advanced breast cancer, which ultimately lead to FDA approval in 1998 (*1*). Since then, targeted cancer therapies have become an accepted therapeutic modality for the treatment of cancer and have contributed to a decrease in cancer related mortality (*2*). However, the benefit of targeted therapies to date has been restricted to 5-10% of solid tumors addicted to oncogenes (3–5). Identifying these relatively rare patients via predictive diagnostic tests relying on genomic biomarkers has created Precision Medicine (*6–8*).

Retrospective analyses of several clinical studies of breast, gastric or lung adenocarcinoma identified increased receptor and/or growth factor expression as prognostic markers for patients with poor prognosis, which highlights the role of ligand-induced signaling as oncogenic drivers (*9–12*). Here we aim to decipher what drives ligand-induced proliferation.

We present the first comprehensive proliferation screen across 58 cell lines comparing to which extent the growth factors EGF (Epidermal Growth Factor), HRG (Heregulin), IGF-1 (Insulin Growth Factor 1) and HGF (Hepatocyte Growth Factor) induce cell proliferation. We find that about half of the cell lines do not respond to any of the ligands whereas the other half of the cell lines respond to a least one ligand. We compare the observed ligand-induced proliferation with the response to treatment with antibodies targeting the ErbB receptor family members, a subfamily of four closely related receptor tyrosine kinases (RTKs): EGFR (ErbB1), HER2/c-neu (ErbB2), HER3 (ErbB3) and HER4 (ErbB4) as well as the Insulin Growth Factor Receptor (IGF-1R) and the Hepatocyte Growth Factor Receptor (Met). Not surprisingly, the antibodies targeting the respective RTK inhibit ligand-induced proliferation. The antibodies also inhibited basal proliferation in some cell lines that do not respond to exogenous ligand addition, which could be driven by autocrine signaling.

The need has been recognized for computational approaches to deal with the complexity of signal transduction and its dysregulation in cancer to ultimately understand drug activity (*13–17*). Large collections of genetic and genomic data lead to efforts to disentangle the complex mechanisms using machine-learning algorithms (18–21). It was previously shown that simulated patient-specific signaling responses derived from mechanistic signaling models using RNA sequencing data from patient biopsies can be robust biomarkers that are predictive of patient outcome (*22*). Here, we combined machine learning and mechanistic modeling to predict which cell lines proliferate in the presence of ligand. We used RNA sequencing data as inputs into a comprehensive mechanistic model capturing the ErbB, IGF1-R and Met signaling pathways. Our novel approach uses simulated signaling features and mutation status of a specific cell line as inputs into a Bagged Decision Tree, which predicts whether tumor cells proliferate in the presence of a growth factor. We achieved a substantial gain in accuracy compared to predictions based on RNA sequencing data alone by inclusion of simulated signaling features such as the area under curve of distinct heterodimers and phosphorylated S6 for *in vitro* models.

Applying this approach to patient data, the prediction of ligand-dependent tumor samples based on mRNA data from The Cancer Genome Atlas (TCGA) revealed that colorectal and lung cancer are the two indications most responsive to EGF, which agrees with the approval of EGFR inhibitors in these indications. In addition, the prediction of responders in patient samples revealed a correlation between predicted tumor growth and measured ligand expression in the tumor microenvironment, which argues for a co-evolution of ligand production and the ability of the tumor cells to respond to stimulation.

## Results

### *In vitro* proliferation screen

To investigate growth factor-induced proliferation we screened a panel of 58 cancer cell lines (10 ovarian cancer, 11 breast cancer, 13 lung cancer, 11 gastric cancer and 23 colorectal cancer cell lines) for response to the exogenously added ligands EGF, HRG, HGF and IGF-1 (Suppl. Fig. 1) that bind to EGFR, ErbB3, Met and IGF-1R, respectively. In addition to ligand stimulation, cells were also treated with ligand blocking antibodies: MM-151, an oligoclonal therapeutic composed of three monoclonal antibodies targeting EGFR (*23*); Seribantumab (MM-121), a monoclonal antibody targeting ErbB3 (*16*); MM-131, a bispecific antibody co-targeting Met and EpCAM (*24*); and Istiratumab (MM-141), a bispecific antibody co-targeting IGF-1R and ErbB3 (*25*). Fig. 1A illustrates the Receptor Tyrosine Kinases (RTKs), their corresponding ligands and the mechanism of action of the ligand blocking antibodies. Proliferation was quantified in a 3D spheroid formation assay at the three-day time point (Fig. 1B) by measuring ATP content as surrogate for cell number (CellTiter-Glo® assay). Response was classified as positive if the signal at the three-day time point was more than 20% above the respective control, plus being significant at a confidence level *α* = 0.05 (measured in quadruplicates, Wilcoxon rank-sum test). Per this screen approximately 45% of cell lines responded to EGF, 55% of cell lines responded to HRG, 33% of cell lines responded to HGF and 7% of cell lines responded to IGF-1. The low response rate to IGF-1 in this proliferation screen may reflect the presence of IGF-1 in the low-serum medium and the modest absolute inhibition point to the importance of IGF-1 mediated signaling for survival rather than for proliferation (*26*). We and others observed a generally weaker MAPK activation via IGF-1R (see Fig. 3C) compared to the other growth factors in the screen (*27, 28*). Further, the CellTiter-Glo® assay relies on metabolic function and hence can be limited as readout for IGF-1 stimulation (*29*).

**Fig. 1:**
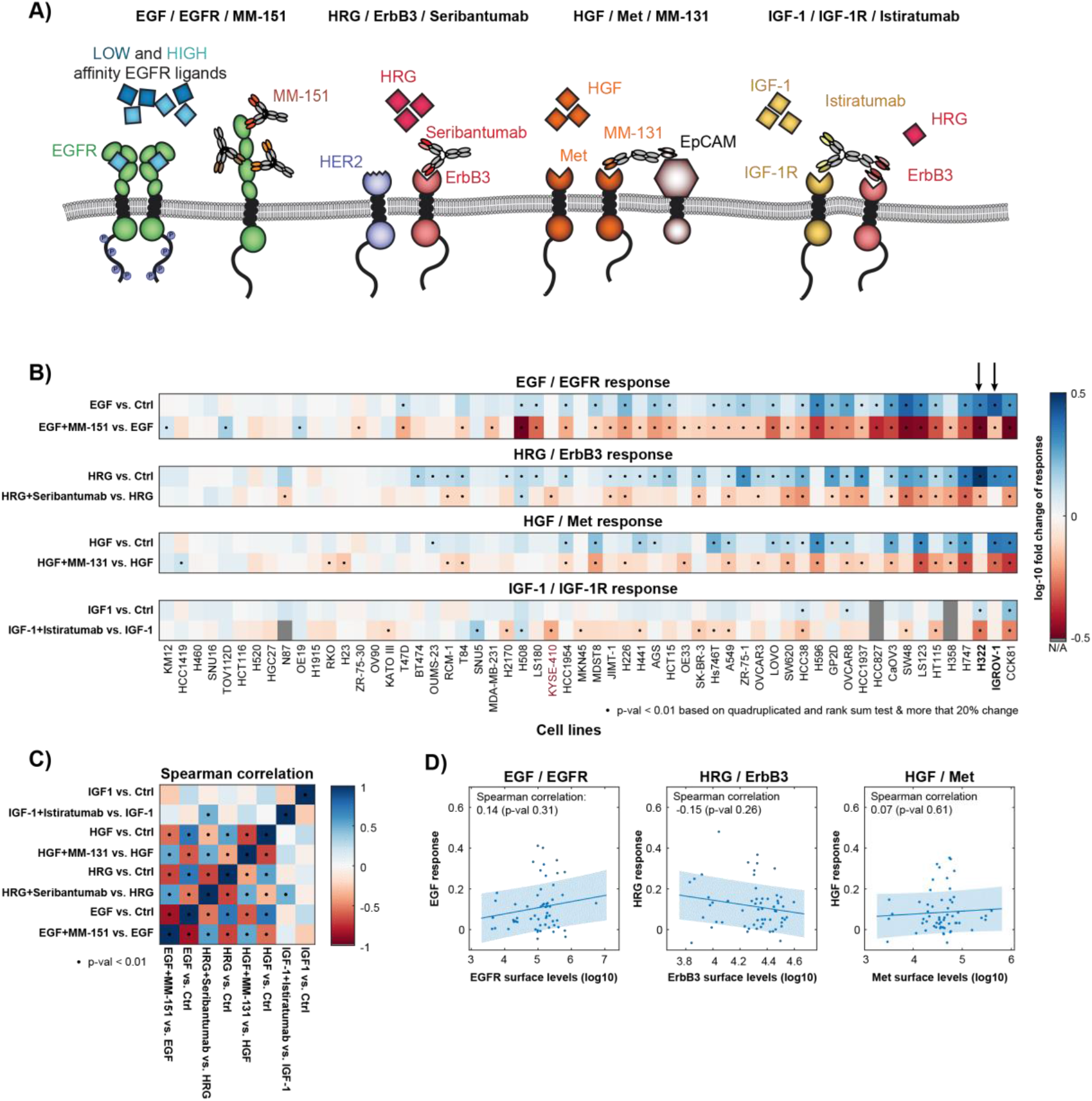
Proliferation screen across 58 cell lines. **A:** Ligand/Receptor and antagonistic antibodies used in the *in vitro* proliferation screen. **B:** Results of the proliferation screen across 58 cell lines. Dots mark a significant increase in ligand induced proliferation or decrease in the presence of ligand plus antibody. The ligand effect is normalized to the medium control, whereas the antibody plus ligand effect is relative to ligand alone. The cell lines marked with an arrow were used to train the computational model to signaling data. **C:** Correlation pattern of ligand and antibody effects across all cell lines. **D:** Linear correlation of receptor expression levels to ligand induced proliferation.

Fig. 1B shows the response to treatment with ligand in combination with the respective blocking antibody compared to the ligand effect alone. Depending on the ligand treatment, 5-17% of cell lines were ligand non-responsive, but the antibodies inhibited basal proliferation, which is indicative of autocrine driven proliferation. Even though IGF-1 did not induce a proliferative response in most cell lines, MM-141 inhibited proliferation in about 19 % of the cell lines indicating that IGF-1 might be present in low-serum medium.

Investigation of correlations between the ligand and antibody responses across all cell lines revealed a checkerboard pattern of significant positive correlations between EGF, HRG and HGF as well as anti-correlations of those ligands and their respective antibody responses (Fig. 1C). This suggests a general trend that cell lines are either responsive to multiple ligands and their respective antibodies (right hand side of Fig. 1B), or are generally non-responsive to any given ligand or antibody (left hand side of Fig. 1B). For IGF-1/IGF-1R, the only significant correlation was observed between Istiratumab treatment and Seribantumab treatment. This can be attributed to both antibodies (co-) targeting ErbB3, which leads to some cell lines responding to both Istiratumab and Seribantumab independent of an IGF-1 effect (see e.g. KYSE-410 cell line in Fig. 1B). The general lack of correlation patterns for IGF-1/IGF-1R responses as were observed for the ErbB family and HGF/Met can be explained by the lack of IGF-1 induced proliferation in this screen.

In the following, we will focus on the question of how ligand dependence can be predicted. A necessary condition for response to any given ligand is the presence of its respective receptor. First, we used a univariate analysis (Fig. 1D) and found that receptor expression levels measured by qFACS do not correlate significantly with the respective ligand response. Therefore, a simple linear model cannot stratify responsiveness. Second, we applied a multivariate approach to predict ligand-dependence. To predict efficiently the phenotypic response and to obtain a comprehensive interpretation of the chosen features, we used a decision tree classification algorithm for the multi-dimensional information about RTK expression levels together with the information about the exogenously added ligand as well as the mutational status of the cell lines (Fig. 4A) (*30, 31*). This resulted in an improvement of the predictions, but it was neither significant nor robust across the ligands studied (Fig. 5A). Next, we investigated whether a multi-pathway signaling model featuring the complex receptor interactions as well as the cross-talk between the mitogen-activated protein (MAP) kinase and the phosphoinositide 3-kinase (PI3K) signaling pathways can be used to predict the phenotypic response. Specifically, signaling features like the area under curve (AUC), quasi steady-state and the signal amplitude of receptor homo -or heterodimers and downstream components were considered as inputs into the decision tree (Fig. 4A).

### Multi-pathway computational model

To construct a comprehensive signal transduction model that could be used to predict proliferation in response to growth factors for all 58 cell lines, we built on a previously published model of ErbB receptor signaling (*16*). We extended the computational model to include IGF1-R and Met (Fig. 2A) as well as twelve homo- and heterodimers for which biological evidence can be found (32–35). Our analysis considers EGFR, HER2, ErbB3, Met and IGF-1R homodimers as well as the heterodimers EGFR-HER2, EGFR-ErbB3, EGFR-Met, HER2-ErbB3, ErbB3-Met, IGF-1R-IGF-1R, EGFR-IGF-1R and HER2-IGF-1R. The latter two were later removed from the computational model without impacting the model performance. Fig. 2B depicts the structure of the model for the example of a signaling HER2-ErbB3 heterodimer. The complete model consists of 62 differential equations and replicates the model structure shown in Fig. 2B for each of the considered ten homo- and heterodimers. In short, receptors bind ligand with published dissociation constants (K_D_). Bound receptors can form homo- and heterodimers and subsequently undergo endocytosis. After internalization, the receptor dimers can get either dephosphorylated and recycled to the cell surface or they get degraded in the lysosomes (*36, 37*). Downstream of the receptor, all homo- and heterodimers except the ErbB3-homodimer, which cannot trans-phosphorylate due to its lack of intrinsic kinase activity, can activate the MAP kinase cascade as well as the PI3K/AKT pathway. ERK and AKT phosphorylation converge in the phosphorylation of S6K1 and S6. Several known feedback mechanisms between the pathways (*27*) were implemented in the computational model. Mathematical details, executable code to simulate the model and instructions to replicate our findings are available in the supplementary material and on biomodels.org.

**Fig. 2:**
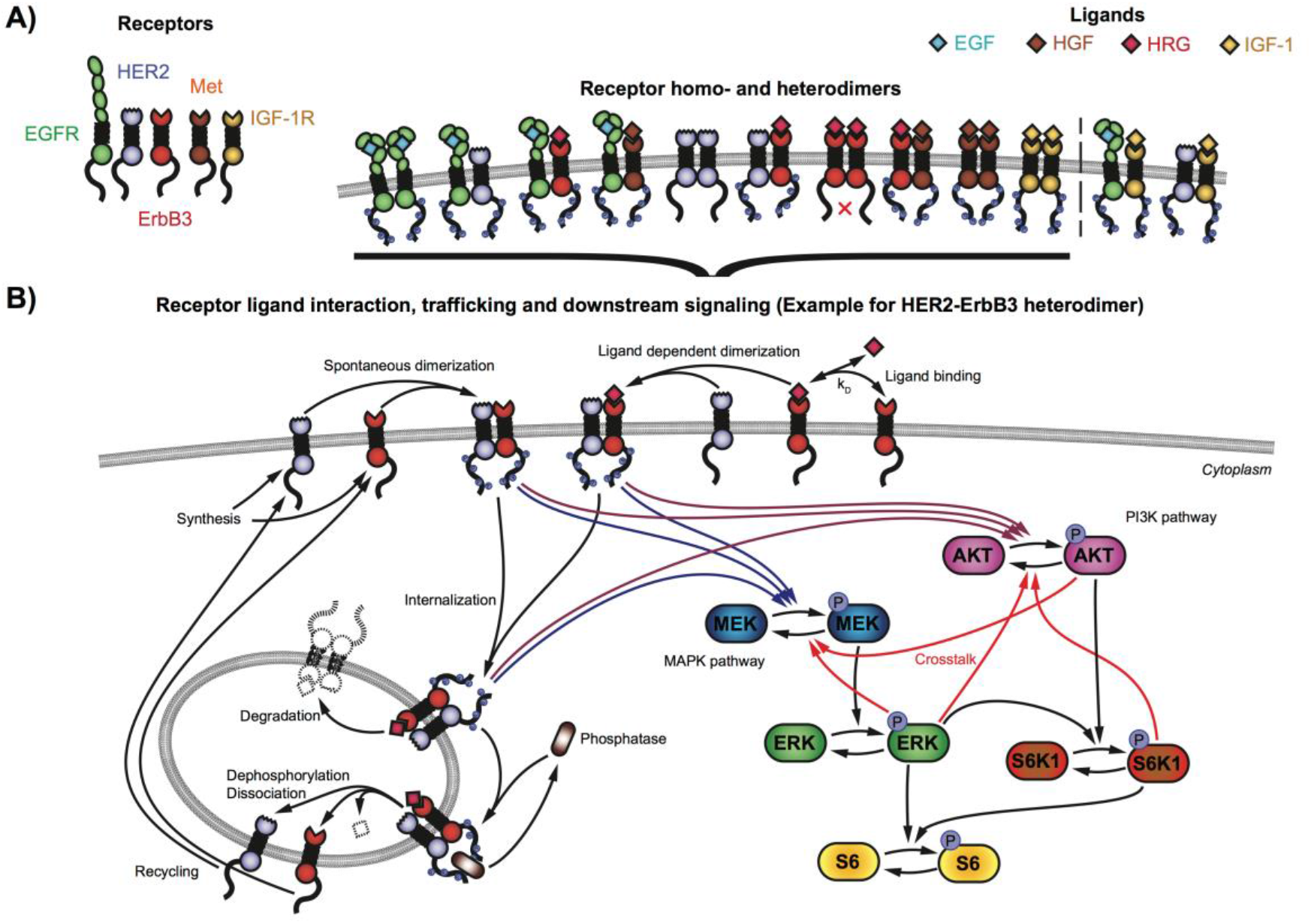
Structure of computational signaling model. **A**: The receptors EGFR, HER2, ErbB3, Met and IGF-1R can form several homo- and heterodimers after ligand binding. **B**: In the model, receptors are synthesized and either dimerize spontaneously or bind a ligand to form homo- and hetero-dimers, which leads to transphosphorylation of the receptors. Activated receptors signal downstream and are prone for internalization, which leads to either degradation or dephosphorylation by a phosphatase followed by recycling to the cell surface. Downstream, the MAPK and PI3K cascade activate S6K1 and ultimately converge in the phosphorylation of S6. The MAPK and PI3K signaling pathways are interconnected via multiple crosstalk mechanisms.

The computational model is constructed with the aim to capture the signaling dynamics of key components of the signaling pathway including receptor homo- and heterodimerization. It is not intended to be a complete compendium of all the known molecular interactions (*35, 38, 39*). Size and complexity of the computational model were chosen to reflect the available experimental data, and to facilitate efficient computation. This is particularly important during model parameter calibration, which uses parameter estimation algorithms to match the available experimental data as closely as possible (see Methods section for details). Mutations were not implemented in the computational signaling model as they appear to increase the signaling baseline but not necessarily the signaling dynamics (*40*). However, the mutation status for each cell line was used in the machine learning classification.

For model calibration, phosphoproteomic time course data from protein microarrays (*41*) for the receptor phosphorylation as well as for phospho-MEK, phospho-ERK, phospho-AKT and phospho-S6 across all seven cancer cell lines (H322M, BxPc-3, A431, BT-20, ACHN, ADRr and IGROV-1) were used. Only the two cell lines H322M and IGFROV-1 were included in the cell line proliferation screen in Fig. 1. These seven cancer cell lines represent different cancer indications (lung adenocarcinoma, pancreatic, epidermoid, breast and ovarian cancer) and were selected based on the molecular diversity with respect to the mutation status and differences in receptor expression levels. A key challenge for building computational models that can describe and predict signaling dynamics of different cell lines is to limit the number of model parameters that are specific to one cell line (*42*). In this case, it was possible to restrict all kinetic rate constants to the same value and to adjust only the receptor expression levels for individual cell lines. Due to the analytically calculated basal activation levels of all homo- and heterodimers as well as of the downstream components, which were derived from steady-state constraints (*43*), the receptor levels impact the signaling response throughout the model. Therefore, the individual receptor expression levels of each cell line enable distinct model responses upon ligand stimulation. The receptor expression levels were measured using quantitative flow-cytometry (qFACS) in combination with RNA sequencing data (see Methods Section and Suppl. Section S4 for details). The model can accurately describe the time course data of seven training cell lines, with 85.7 % of the data points within two standard deviations of the model uncertainty (see Fig. 3 for a selection of the data and Suppl. Figs. 12-41 for a comprehensive comparison of model simulations and experimental data).

**Fig. 3:**
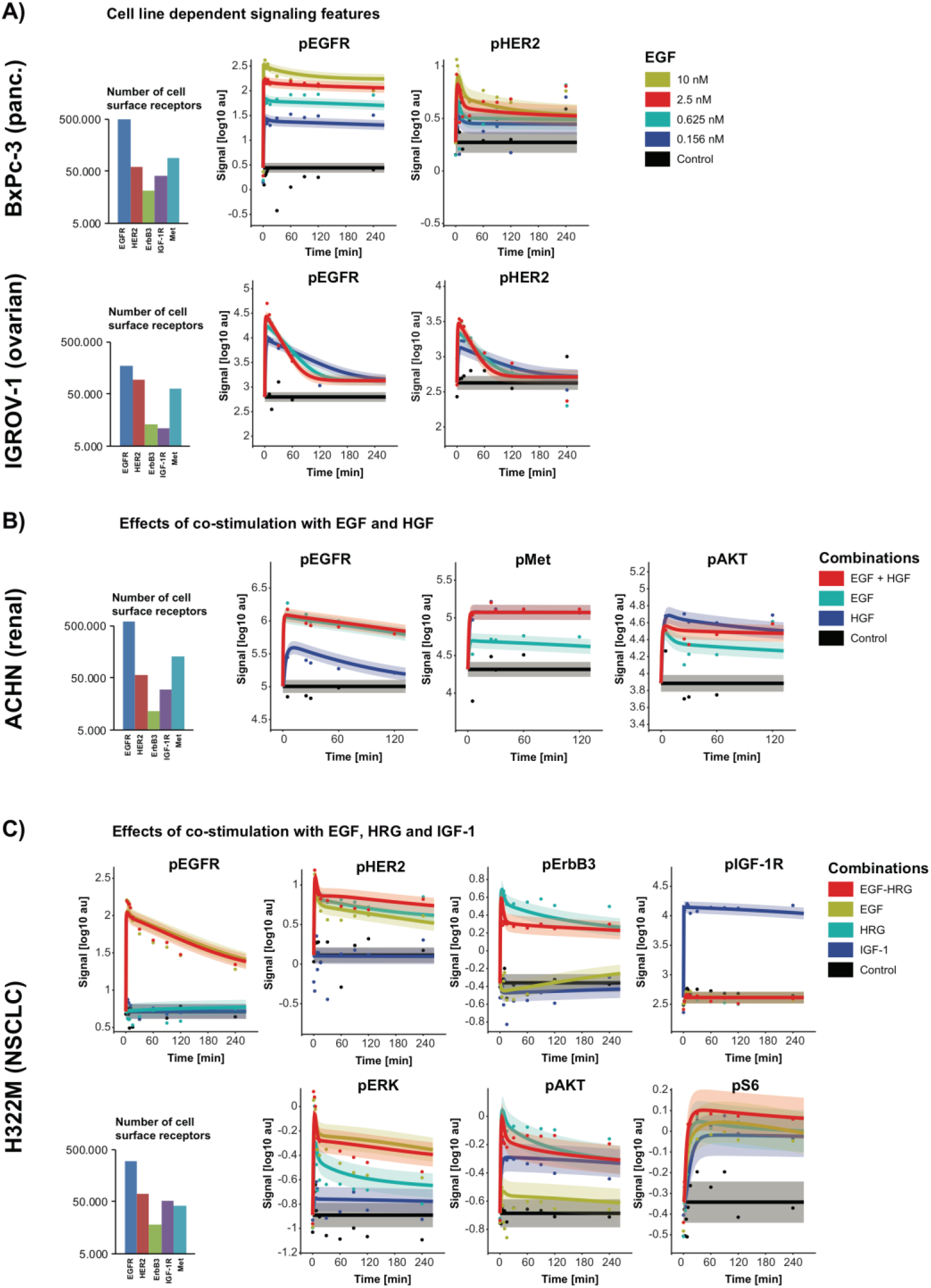
Importance of receptor surface levels for model response, shown for a selection of calibration cell lines. **A:** Cell line dependent signaling features: Model response to EGF stimulation of two different cell lines leading to sustained or transient receptor phosphorylation in the BxPc-3 and IGROV-1 cells. Their respective receptor surface levels are shown on the left. The model fits are represented by the colored lines with respective uncertainties (67% confidence intervals) as shades. Data points are shown as dots in the same color. **B:** Model fits for the cell line ACHN stimulated with HGF, EGF and the combination. C: Model response to co-stimulation of EGF plus HRG in comparison to the stimulation with EGF, HRG or IGF-1 alone in H322M cells.

### Validation of the computational signaling model

Based on the trained model, predictions were generated for two independent validation cell lines (BT-474 M3, MDA-MB-231) and compared to the experimental data. The goodness of the predictions for the validation cell lines was equivalent to the goodness of fit of the training cell lines (Fig. 3B, Suppl. Figs. 36-41 for model fits to the available data). These simulation results validate that receptor expression levels are sufficient to predict signaling features of independent cell lines that were not used for model training. In addition, we generated model predictions that were based on random receptor surface levels, by taking non-matching values from randomly selected cell lines used in the cell viability assay (see **Error! Reference source not found.**). The decline in goodness of fit was on average 30% and statistically significant (*p* = 8.5 * 10^−9^, see Suppl. Fig. 2). These results illustrate the importance of receptor expression levels and their ratios to capture the distinct signaling features observed for each cell line (*44*).

To further validate the presented model structure with its multiple receptor heterodimers, a simplified model lacking any heterodimerization capabilities was trained to the experimental data. In this setting, all receptors could signal downstream through homodimerization, disregarding the non-functional kinase unit of the ErbB3 receptor. Even with 39 parameters less, the reduced model had a goodness of fit impediment with associated p-value < 1.e-15 in the corresponding likelihood-ratio test, showing the significant improvement of the computational model by including receptor heterodimerization. Besides, concordance of the basal receptor levels obtained via analytic steady state equations, reflecting the proposed receptor trafficking, was assessed through extensive measurements of basal total and phosphorylation levels in 39 breast cancer cell lines (*28*). A good correlation, especially for the ErbB receptor family, was found (see Suppl. Fig. 3), confirming the calibrated model parameters constituting cell-dependent steady states.

To further test the robustness and applicability of the model to ligands not included in the original training-set, we compared the predicted receptor activation patterns in response to different ligands of the EGFR-ligand family, such as Betacellulin (BTC). To this end, previously published (*16*) time-resolved data of the ADRr cell line for EGF and BTC with ligand concentration range between 0.1nM and 10nM was reanalyzed with the current model (Suppl. Fig. 4A). Differences in the ligand binding affinities of each ligand to the EGF receptor as well as different homo- as well as heterodimerization kinetics were sufficient to describe the experimental data (Suppl. Fig. 4B and C). BTC induces a stronger EGFR homodimerization compared to the stronger EGFR-HER2 heterodimerization induced by EGF (Suppl. Fig. 4D). These differences in EGFR homo- and heterodimerization with HER2 were previously described (*45, 46*).

### Importance of receptor homo- and heterodimers

Further insights into growth factor signaling and signal processing by the cancer cells can be gained by analyzing the computational model and why it can capture the distinct signaling dynamics across cell lines. This analysis revealed the importance of different homo- and heterodimers in encoding information as a function of the ligand(s) present. The largest effect of heterodimerization on signal output is seen within the ErbB family. The interplay between the receptors explains the slower, more sustained receptor activation in response to EGF in the BxPc-3 cells, which are characterized by a high ratio of EGFR to other receptors (Fig. 3A). In contrast to the BxPc-3 cells, the IGROV-1 cells are characterized by low EGFR expression levels compared to other receptors leading to the observed transient and early activation of EGFR and HER2. For the ACHN cell line, we generated signaling data in response to EGF, HGF as well as to the combination of EGF and HGF. The computational model reveals sophisticated feedback regulation between the MAPK and PI3K pathways, e.g. reduced AKT activation comparing EGF and HGF co-stimulation to HGF only (Fig. 3B). In Figure 3C another example is depicted: when EGF and HRG are present, EGFR and ErbB3 compete for HER2. The ligand combination leads to reduced phopsho-ErbB3 levels due to a dominant binding of HER2 to EGFR in the presence of EGF (Fig. 3C). We argue that mechanistic understanding of changes in receptor stoichiometry based on individual ligands or ligand mixtures that lead to non-obvious signaling responses is required to understand the ultimate phenotypic response.

IGF-1 is distinct from the other growth factors in our screen. IGF-1 displayed a much weaker ability to induce proliferation and similarly the effect of IGF-1 signaling in co-stimulation experiments is distinct to the HRG/EGF or HGF/EGF co-stimulation experiments. The time-course data for co-stimulation of IGF-1 with either EGF or HRG did not show deviations from the respective stimulation with IGF-1 alone (see Suppl. Fig. 15). Consequently, the model parameters referring to the heterodimerization of IGF-1R (see heterodimers involving IGF-1R in Fig. 2A) with other receptors could be set to zero without a significant decline in the goodness of fit.

### RNAseq and signaling features are predictive of phenotype

The calibrated mechanistic signaling model can be used to simulate signaling features for the stimulation with EGF, HRG, HGF or IGF-1 for all cell lines of the cell viability screen only using their receptor expression levels as inputs. To connect signaling features derived from the computational model to the phenotypic response observed in the cell proliferation screen, we applied a machine learning approach. Based on different sets of input features that were selected based on their prediction ability (see below), we trained bootstrap-aggregating (bagged) decision trees (BDTs). BDTs are highly efficient for multivariate analysis and allow for a comprehensive interpretation of the chosen features (*47, 48*). Therein, a multitude of trees are trained, with each single tree aiming at discriminating growing from non-growing cells based on the provided feature space. At each node, a tree divides the data through selected features that yield the best improvement in signal-to-noise. Combining the ensemble of trees, high-dimensional and non-linear regions in feature space prone for cell growth are obtained and can be used for prediction. More details can be found in Suppl. Section S2.

To investigate the contribution of dynamic signaling features derived from the computational model on the predictive performance of the machine learning approach, we considered two different sets of input features (additional sets are reported in Suppl. Fig. 5). The first feature set contained the receptor expression levels, KRAS and PI3K mutation status and ligand treatment as binary input (Fig. 4B). The second feature set did not contain the receptor expression levels explicitly but only signaling features derived from the computational model based on the receptor expression levels and the ligand stimulation (Fig. 4C) as well as the mutation status (see **Error! Reference source not found.**). The cell line specific signaling features consist of the area under the curve (AUC) of all phosphorylated receptor homo- and heterodimers in addition to AKT, ERK and S6 phosphorylation. Inclusion of the fold-change of these features as well as their quasi steady-state levels were tested but did not yield substantial benefits on top of the information given by the area under curve (see Suppl. Fig. 5). Fig. 4 illustrates both multivariate prediction strategies as well as the univariate analysis shown in Fig. 1D.

**Fig. 4:**
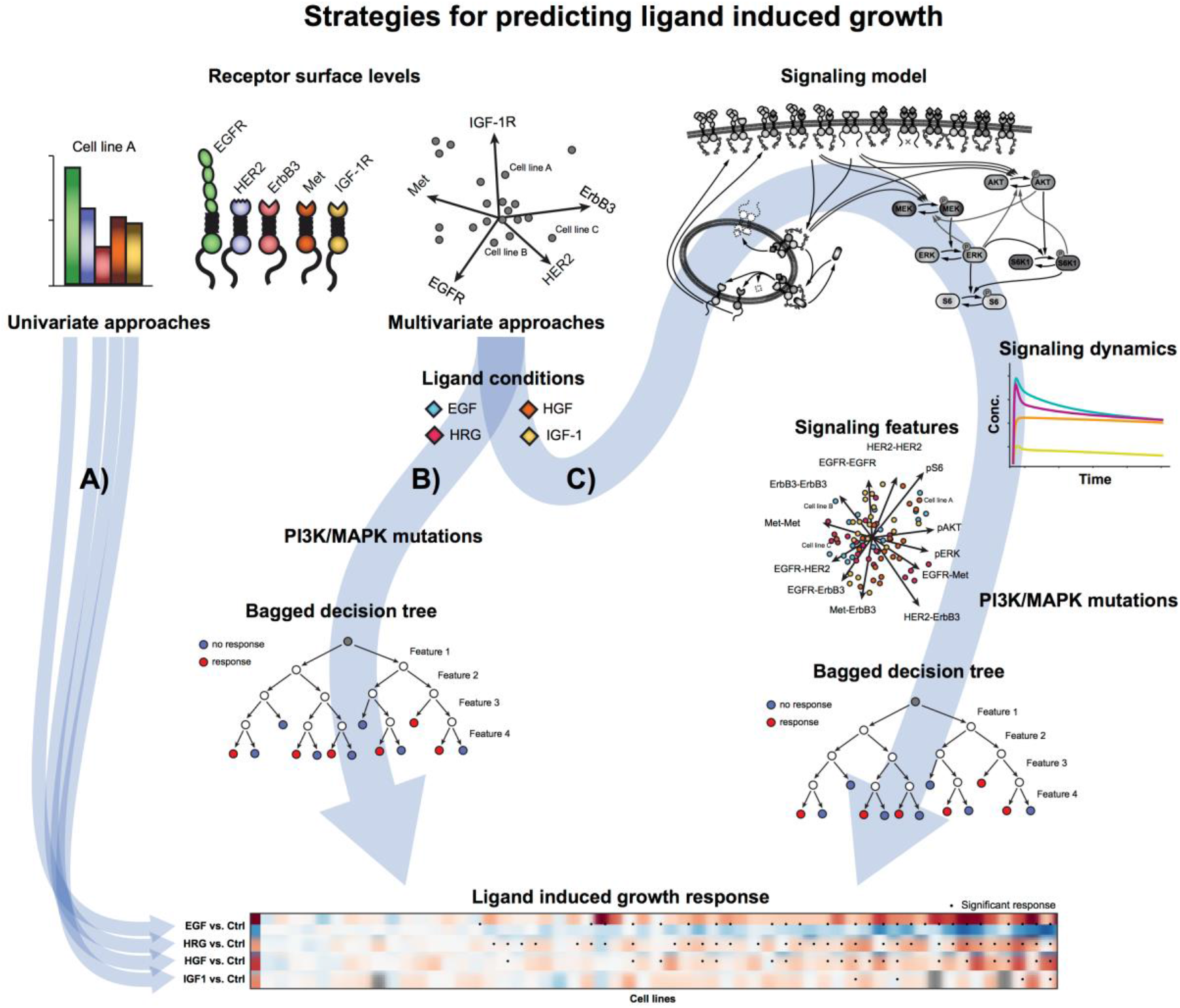
Strategies for predicting ligand-induced phenotypic response. Based on the receptor expression levels of individual cancer cell lines, either a univariate or multivariate approach can be used to predict the phenotypic response to ligand stimulation. **A:** Univariate approaches relate the respective receptor levels to the observed ligand induced proliferation for each of the four ligands separately. **B-C:** Multivariate approaches such as bagged decision trees (BDTs) relate high-dimensional feature sets to the observed phenotype. **B:** In this case, the feature set consists of the five receptor surface levels as well as information about the respective ligand stimulation and mutation status. **C:** The calibrated and validated signaling model allows to simulate the expected signaling dynamics for each individual cell line based on its receptor expression levels and ligands present. Based on the mechanistic knowledge that the signaling model incorporates, it can expand the initial five-dimensional feature set to a 12-dimensional feature set. This expanded feature set, together with information about mutation status is now connected to the observed growth responses by a bagged decision tree.

To evaluate the accuracy of bagged decision tree (BDT) predictions, the cell lines were randomly split 500 times into training and testing sets. For each ligand, BDT training was performed on the training data for all available ligands, while efficiency was calculated on the testing cell lines for the chosen ligand only. By leaving out whole cell lines as opposed to a fraction of the total data, possible bias due to correlated responses to different ligands in the same cell line is avoided. We monitored the fraction of true predictions as a metric for the prediction accuracy. Both feature sets resulted in a better prediction of proliferation compared to random data, which results in 50% true predictions and serves as control (Fig. 5A). Exceptions were the predictions for IGF-1 and HGF stimulation using the receptor expression levels only, where the performance drops insignificantly below the control. Training on features derived from the computational model improved the prediction of cell proliferation significantly compared to control (p-value of <1.e-2, see Fig. 5B) for the combination of all ligands except IGF-1, while BDT predictions based on receptor expression levels alone did not result in a statistically significant improvement (p-value of 0.15, see Fig. 5B). The respective distributions for single ligand induced proliferation predictions can be found in Suppl. Fig. 6. The BDT training was robust with respect to the relative amount of training and testing data and to the significance threshold upon which a cell is labeled as proliferating (Suppl. Fig. 7). For IGF-1, the low number of responders resulted in a low correlation within the events and a statistical bias of their relative amount in training or testing data. These circumstances rendered robust prediction impossible and the IGF-1 data set was excluded, e.g. in the combination of all ligands (see Fig. 5A).

**Fig. 5:**
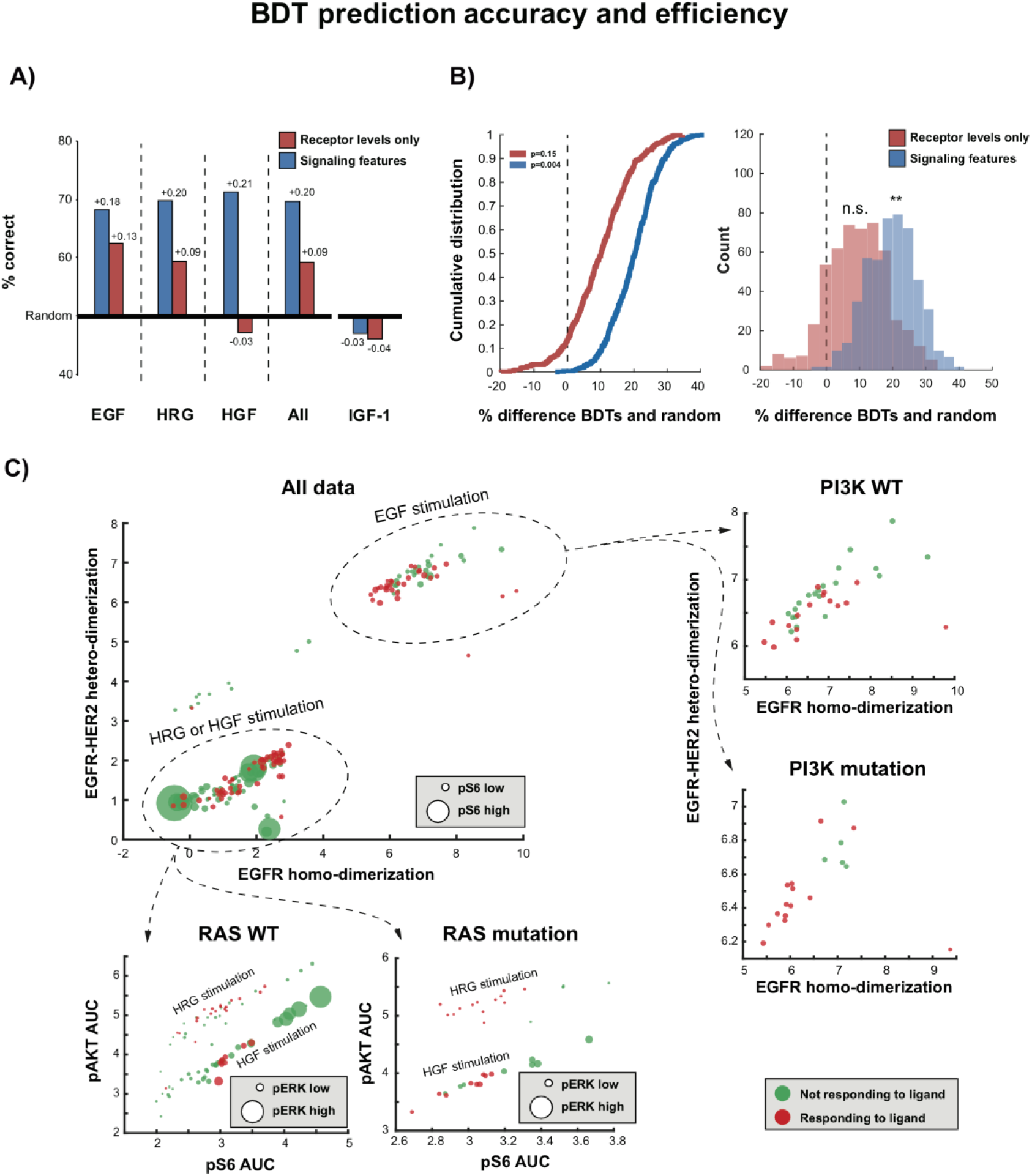
Prediction of ligand-induced proliferation using BDTs. **A:** Ratio of true predictions after BDT training with simulated signaling features or receptor expression levels only, compared to random predictions in the presence of EGF, HRG, IGF or HGF. **B:** For 500 random splits of training and testing cell lines, the BDT outcome is compared to random growth assessment as histogram and cumulative density function, showing the significant improvement due to mechanistic modeling. **C:** Data of *in-vitro* cell viability screen that leads to either proliferation (red) or did not alter the phenotypic response significantly (green) in different 2D representations of the feature space.

One of the advantages of mechanistic computational models is that it is possible to gain insights into cellular signal processing. Thus, the importance of different model features during BDT training can be traced back to develop hypotheses about what ultimately drives proliferation. The training features are ranked by their impact on data classification, which is measured by the average gain in signal-to-noise ratio over all trees. As illustrated in Table 1, the most important features rely on the homo- and heterodimerization stoichiometry of the ErbB receptor family as well as on the downstream signaling. A more detailed overview is given in Fig. 5C, which illustrates the data from the proliferation screen and the model-derived features that proved important during training of the BDT (see Table 1). This coincides with prior biological knowledge that ErbB receptors induce proliferation (*13*). It can be observed that EGF, HGF or HRG stimulation induce very specific homo- and heterodimerization patterns of EGFR and EGFR/HER2 respectively. Moreover, phospho-S6 is an important feature to predict proliferation in the presence of HRG and HGF, both mainly activating the PI3K pathway. Its importance might be a result of the crosstalk between the MAPK and PI3K pathways and the convergence of both pathways in S6 phosphorylation (*49*). Moreover, PI3K mutation status and EGFR heterodimerization patterns help to identify clusters of EGF dependent cell lines. RAS mutation status and differences in activation of downstream targets further predict proliferation after HRG and HGF stimulation. Independent of the KRAS mutation status, HRG induces increased levels of AKT phosphorylation while inducing the same amount of S6 phosphorylation as HGF.

**Table 1:** Model features ranked by their BDT training efficiency.

| Feature w IGF1, HRG/IGF1 | BDT importance score |
| --- | --- |
| pS6 AUC | 0.65 |
| EGFR homodimerization | 0.56 |
| EGFR-HER2 heterodimerization | 0.56 |
| Met-ErbB3 heterodimerization | 0.55 |
| pAKT AUC | 0.43 |
| pERK AUC | 0.37 |
| PI3K mutation status | 0.35 |
| RAS mutation status | 0.30 |

### Application to patient data

In the previous section, we illustrated the power of mechanistic computational models to predict *in vitro* proliferation. Next, we applied this novel approach to predict ligand-dependent proliferation to patient derived tumor samples. Specifically, data generated by the TCGA Research Network (http://cancergenome.nih.gov/) was utilized, and ligand-dependent, proliferating tumors were predicted using the signaling model and machine learning algorithm outlined in the previous sections. The data set includes 2909 samples from patients with breast, colorectal, lung and ovarian cancer. As the measured RNA expression levels between cancer cell lines and solid tumor tissue were on different scales, we normalized the expression levels to the mean. Subsequently, the expression levels of EGFR, Her2, ErbB3, Met and IGF-1R were used to perform model simulations and to extract the signaling features needed to predict ligand-dependent tumors using the previously described BDT algorithm. The number of ligand-dependent tumors differed within indications and ligand (EGF, HGF or HRG). The number of predicted ligand responsive tumors was highest for HRG followed by EGF and lowest for HGF (Fig. 6A). Lung and colorectal cancer seem to be most responsive to EGF, which is congruent with the high prevalence of EGFR mutations and overexpression in these indications (50–52). In Fig. 1B we observed that ligand induced proliferation is correlated with treatment response to an antibody targeting the respective receptor. Therefore, the approvals of EGFR inhibitors in non-small cell lung cancer (NSCLC) and CRC confirm the predicted dependence on EGFR signaling (*53, 54*). In contrast, the low dependence of NSCLC on HGF signaling might explain the failure of Onartuzumab (MetMAb), a Met blocking antibody in a Phase 3 study in NSCLC (*55*). Similarly, EGFR inhibitors have not yet proven to lead to clinical benefit in breast cancer (*56*) which is also in agreement with the predicted low EGF dependence of breast cancer. The predicted high responsiveness of breast cancer and lung cancer to HRG seems to agree with the retrospective analysis of two clinical studies with Seribantumab and the finding that HRG expression appears to be predictive of patients responding to therapy (*57*).

**Fig. 6:**
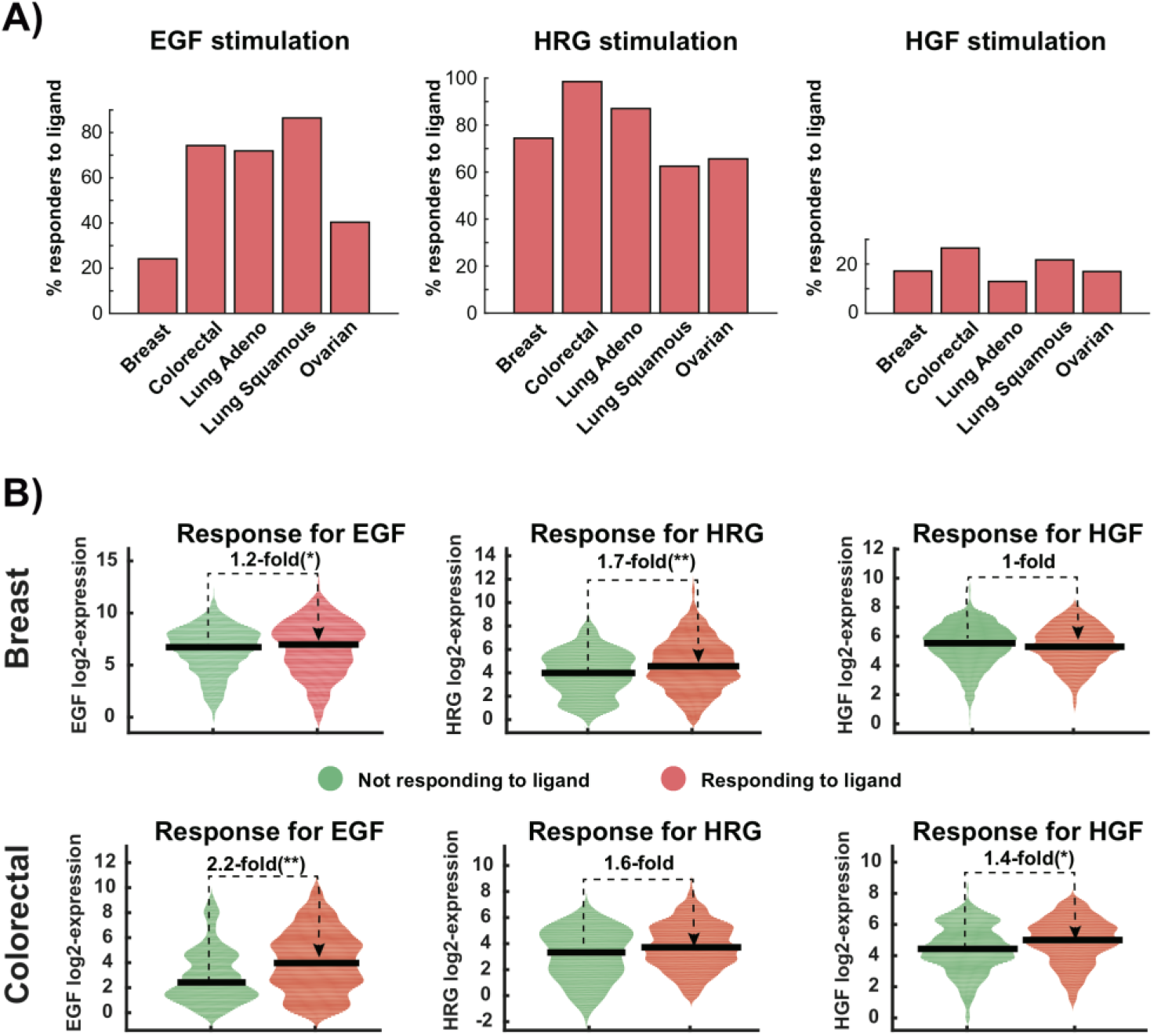
Predicting ligand dependent tumors from the TCGA data set. **A:** Patient-derived expression levels for breast, colorectal, lung and ovarian cancer were utilized to classify tumors into ligand-dependent and ligand-independent tumors **B:** The measured RNA expression of the respective ligands is plotted for ligand-dependent and ligand-independent tumors, the line indicates the mean expression and the stars indicate a statistically significance of difference.

The predicted ligand-dependence only considers the molecular makeup of the tumor and therefore only indicates if a tumor would respond to ligand if present in sufficient amounts. The local concentration of the ligands, however, cannot be inferred from our analysis and it is difficult to match to the *in vitro* data that was used for model training. This uncertainty impacts the predicted number of ligand-dependent tumors in this data set. Apart from the relative number of ligand-dependent tumors, we observed a significant correlation between ligand expression levels and the predicted response to ligands (Fig. 6B, additional data in Suppl. Fig. 8). The predicted ligand-dependent tumor samples from patients with breast and colorectal cancer display statistically significant higher (t-test) amounts of the corresponding ligand compared to the predicted ligand independent tumors, if we compare the mean expression levels. This suggests that tumors that express ligands evolved to utilize them, and vice versa.

## Discussion

The research to understand and therapeutically battle various cancer types has made significant progress over the last two decades, decreasing the overall mortality by roughly 2 % per year since 2001 (*58*). Targeted therapy and combinations thereof have become important areas of drug development with rising FDA approval rates in the last years (*59, 60*). However, by studying cellular fate across multiple cell lines and indications, we and others have learned that complex interactions between receptors as well as positive and negative feedback regulation between signaling pathways can diminish drug efficacy. To obtain a deeper understanding of cell response to exogenous stimuli, phenotypic responses need to be studied in the context of multiple signaling pathways as well as mutation status. In this work, we developed a computational model describing multiple signaling pathways and show that a BDT algorithm using simulated signaling features can accurately predict ligand-dependent proliferation *in vitro*.

The signaling model incorporates the ErbB receptor family as well as the Met and IGF-1R receptors. Parameters of the model were estimated based on a variety of time-resolved data from seven different cell lines including a wide range of ligand concentrations with comprehensive single ligand and co-stimulations. The cell lines cover a broad range of ratios of receptor expression levels. The ligand concentrations used in the proliferation screen were in the range of concentrations used for the signaling experiments. While retaining a good fit to the experimental data, we could keep all kinetic parameters in the signaling model constant and just vary the receptor expression levels to describe the experimental signaling data for each cell line. The computational model not only describes the data for the seven training cell lines but also predicts the signaling responses of two additional, independent validation cell lines. We showed that the goodness of fit is dependent on the absolute receptor expression levels and the formation of different homo- and heterodimers. The observed variability in model response was achieved by differences in internalization, degradation and recycling rates for different receptor homo- and heterodimers. Saturation of different downstream model components is sensitive to the receptor expression levels. The analytically solved steady states for all cell lines were important to implement the complex receptor dimerization properties in the signaling model. Thus, ligand stimulation lead to a complex re-distribution of receptor homo- or heterodimerization and facilitated distinct downstream activation patterns. The calculated steady states, especially for the ErbB receptor family, were found in concordance with measured basal total and phosphorylation levels of 39 breast cancer cell lines utilized from (*28*). The fact that the developed computational model can accurately describe the signaling responses across multiple cell lines enables the prediction of signaling dynamics for cell lines of the proliferation screen that were not used to construct the computational signaling model.

To predict proliferation in response to ligand stimulation, we linked simulated signaling features to the data of the cell proliferation screen using a supervised machine learning approach. Tree-based classification algorithms are widely used for machine learning (*61–63*) and carve out regions in feature space that best distinguish between different data classifications, here proliferation vs. stasis upon ligand stimulation. Bagged decision trees (BDTs) were trained on the *in vitro* proliferation screen across 58 cell lines. The algorithm was trained with either receptor expression levels and ligand stimulation conditions as Boolean columns or with signaling features extracted from the computational model. The signaling features included the integrated area under curve of all receptor complexes as well as the phosphorylation of AKT, ERK and S6. Both feature sets lead to a better prediction of proliferation compared to control, where response was predicted based on random data. However, the signaling features allowed for a more robust and statistically improved prediction of proliferation. In addition, the computational model allows us to gain insights into the underlying processes driving ligand-dependent proliferation. In all cases homo- or heterodimerization of the ErbB receptors was important. In the case of EGF, the PI3K mutation status mattered in addition. For HRG and HGF possible RAS mutations together with AKT and S6 phosphorylation were important features to predict cell proliferation. However, the importance of features in the tree-based approach is very sensitive to the utilized data, and additional measurements are needed to infer the role of MAPK and PI3K signaling in inducing growth.

Simulated signaling features are advantageous over using receptor expression levels directly as input features for two reasons: First, the dynamic range of receptor activation as well as of the downstream components is described quantitatively and renders the model outputs more robust to receptor expressions levels, which span multiple orders of magnitude. Second, the interplay between receptors and the included feedback mechanisms adds a source of information on top of the receptor expression and ligand information alone, leading to a non-linear input transformation that improves the detection of regions governing proliferation.

To demonstrate the applicability of this novel approach to patient samples, data from 2909 patients with breast, colorectal, lung or ovarian cancer were analyzed. For these samples, model simulations were conducted to extract signaling features required for BDT prediction of ligand-dependence. Interestingly, we observed a significant correlation between the measured ligand expression levels and the predicted ligand-dependence for breast and colorectal cancer. This may be a consequence of evolutionary adaption of these tumor cells due to the growth advantage from ligand-mediated signaling. Therefore, the presence of ligands in the tumor micro-environment may be a favorable biomarker for RTK-directed drug treatment. This rationale together with possible mutations, which contributed substantially in the BDT training, are currently being explored in the clinic (*57*).

However, both the prediction of proliferation and the mechanistic computational model have limitations. For one, machine learning in feature-space regions that are not covered by many cell lines is not efficient as illustrated by IGF1 in the proliferation screen data-set. The mechanistic model is limited in its ability to fully reproduce data in cases of either receptor overexpression (see Suppl. Fig. 20), which probably transfers to the presence of activating mutations, as well as in cell lines harboring PI3K or RAS mutations (see Suppl. Figs 23, 24, and 38 to 41). We also encountered computational limitations since the complexity of the mechanistic computational model is on the verge of what is currently computationally feasible. This said, more emphasis on model selection, e.g. profile likelihood-based model reduction (*64*) with additional prior knowledge may allow us to better bridge between quantitative time-resolved data and large-scale genomic and phenotypic data. To understand ligand mixtures and pathway redundancy a greater variety of single ligands, e.g. FGF and PDGF, or stimulations with ligand mixtures might aid to more accurately determine model parameters for receptor dimerization, trafficking and downstream activation in the future. Additional perturbations like specific gene knockdown etc. will help to improve the signaling model presented here.

Ligand-dependence or addiction to growth factors (ligands) might prove to be far more prevalent than oncogene addiction given the high ligand prevalence in solid tumors and potentially as much more complex since multiple ligands are expressed in the tumor microenvironment. To successfully treat patients with ligand-dependent tumors with targeted inhibitors (small molecules or monoclonal antibodies) or rational combinations of targeted inhibitors, a better understanding of ligand-dependence is crucial, especially given the redundancy of signaling pathways within a tumor cell and tumor heterogeneity. We argue that the mechanistic understanding of changes in receptor stoichiometry based on individual ligands or ligand mixtures lead to non-obvious signaling responses relevant to the ultimate phenotypic response. The work presented here demonstrates that for targeted therapies to be successful in the clinic the ligand hierarchy as well as co-dependence need to be understood and most likely require the measurement of multiple ligands and respective receptor expression levels in tumor biopsies. New methodologies like single cell RNAseq (*65*) may allow us in the future to characterize the clonal composition of tumors and to determine which cellular fraction is ligand-dependent and which drug combination is best suited to eliminate all ligand-dependent tumor cells.

The presented novel approach of using BDTs in conjunction with simulated signaling features is the beginning of how complex mechanistic models and large data sets can be combined to understand cell-specific complexity but also heterogeneous tumors better. We demonstrated that mechanistic computational models of signaling pathways can help bridge between large scale *in vitro* observations and clinical hypotheses. In the future, selected *in vivo* studies should be used to validate rational combination regimens. Previous efforts predicting drug sensitivity based on large and diverse data sets found that gene expression data proved most valuable, together with exploiting non-linear relationships and addition of prior knowledge of biological pathways (18, 66, 67). Yet, significant improvement in predictions proved to be challenging across multiple approaches and data sets. With the presented approach, mechanistic knowledge can be easily combined with known datasets from RNA sequencing, copy-number alterations and mutation information to improve the prediction of patient-individual drug response and unravel the interplay between complex signaling and cellular fate.

## Materials & Methods

### Experimental time-course data and selected cell lines

The available data consists of three different data sets, which include time-resolved concentration measurements of activated and total receptors as well as of various phosphorylated downstream targets. The largest data set comprises nine cell lines with measurements of the EGFR, HER2 and ErbB3 receptors of the ErbB family and the IGF1-receptor, together with the downstream targets ERK, AKT, S6K1 and S6. Four different ligand concentrations of Epithelial growth factor (EGF), Heregulin (HRG), and Insulin-like growth factor (IGF-1) ranging from 0.156 to 10 nM are used and 12 measurement time points up to 240 minutes are taken. In addition, co-stimulations of the respective ligands are available for two of the nine cell lines. Out of the nine cell lines, six cell lines are used for model calibration, the remaining three to validate the model. Cell lines used for calibration include H322M (non-small cell lung cancer), BxPc-3 (pancreatic cancer), A431 (epidermoid cancer), BT-20 (breast cancer), ADRr (ovarian cancer) and IGROV-1 (ovarian cancer). BT474 (breast cancer), MDA-MB-231 (breast cancer) and ACHN (renal cancer) are utilized to validate the model.

Measurements after either EGF or Betacellulin (BTC) stimulation are available for one of the calibration cell lines, ADRr, with ligand concentrations between 0.11 and 9.26 nM, and 12 measurement time points up to 240 minutes. These include phosphorylation of EGFR, HER2, ErbB3, ERK and AKT. Apart from that, measurements with HGF and EGF as well as their co-stimulation is available for ACHN, which is used for validation with respect to HRG and IGF-1, spanning the same concentrations and measurements up to 120 minutes. Therein, phosphorylated EGFR and Met phosphorylation as well as phospho-ERK and phospho-AKT are measured. The receptor concentrations in all experiments are measured by ELISA whereas the downstream components are measured by lysate microarray (see Suppl. Section S3).

### Mechanistic modeling

Mechanistic models based on ordinary differential equations (ODEs) are frequently used for the description of biochemical reaction networks. They are composed of kinetic rate equations and every component x of the model has a biological counterpart. The time evolution *x(t*) of the model concentrations is obtained by integration of the corresponding system of ODEs

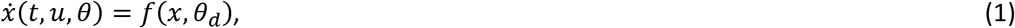

depending on initial and kinetic rate parameters comprised in *θ_d_*. These are linked to measured concentrations of the involved constituents *y*(*t*) by an observational function

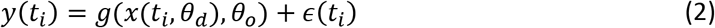

with the assumption of Gaussian errors *ϵ* ~ *N*(0, *σ*) that is often achieved via log transformation. In addition, the observation function includes e.g. scaling and offset parameters, summarized in *θ*_o_. Both observational and dynamic parameters are comprised in *θ*. To compare the model response to measured data at time points *t_i_*, the scaled log-likelihood is calculated via

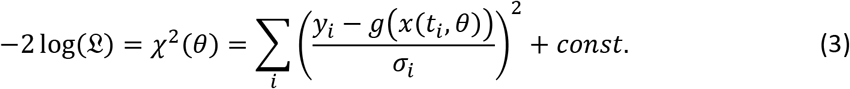

Within the maximum likelihood framework, the optimized parameter set 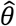 is estimated through minimization of *χ*^2^(*θ*).

Since analytical solutions of non-linear ODE systems are in general not available, a numerical integration must be performed. In this work, the dynamical system and its sensitivities were integrated by the CVODES integrator of the SUNDIALS suite (*68*). Therein, an implicit BDF integration method (*69*) with attached KLU sparse solver was chosen (*70*). The inner derivatives of the likelihood needed in gradient-based parameter estimation were computed via forward sensitivities supplied to the integration algorithm (*71*). Numerical optimization was conducted using a trust-region based, large scale nonlinear optimization algorithm implemented in the MATLAB function LSQNONLIN (*72*). For the mathematical modelling and visualization, the open-source and freely available d2d framework (*73*), based on MATLAB, was used.

### Calculation of receptor surface levels

To calculate receptor surface levels, mRNA expression levels of the receptors utilized in the signaling model were acquired from the Cancer Cell Line Encyclopedia (CCLE) (*74*). Further, they were compared to 124 cell lines, for which qFACS receptor/surface measurements (see Suppl. Section S4) were available at Merrimack Pharmaceuticals (Suppl. Fig. 9). A functional fit to the data was used to calculate receptor/surface expression levels for all cell lines examined in this study (see **Error! Reference source not found.** and **Error! Reference source not found.**).

A good correlation between mRNA and protein expression was previously shown in an independent study (*75*). Receptor surface levels for cell lines that were not included in the CCLE database were taken from in-house qFACS measurements.

## Acknowledgements

We thank Tim Heinemann, Jeffrey Kearns, Sergio Iadevaia, Yasmin Hashambhoy-Ramsay and Tim Maiwald for their constructive feedback and proof reading the manuscript.

## Author contributions

HH build the mechanistic signaling model, built BDT and TCGA predictions, performed the computational analysis and wrote manuscript. KM performed the cell viability screen and wrote the manuscript. MS and JA performed the ELISA and lysate microarray assays. VP performed the qFACS assay. SW helped built the mechanistic signaling model. JT and JS revised the manuscript. BS and GM helped plan the study and wrote the manuscript. AR planned the study, supervised computational work and wrote the manuscript.

